# Developmental switch dichotomizes kidney response to *Nphp3* inactivation and treatment outcome

**DOI:** 10.64898/2026.05.21.726570

**Authors:** Joran Martin, Alice S Serafin, Florine Chereau, Younes Achouri, Nicolas Cagnard, Marie-Christine Verpont, Alexandre Benmerah, Isabelle Scheers, Patrick Jacquemin, Sophie Saunier, Amandine Viau

**Author notes:** Corresponding author. Amandine Viau, Inserm U1163 - Institut Imagine Laboratory of Hereditary Kidney Diseases 24 Boulevard du Montparnasse, 75015 Paris France. Co-first author (Joran Martin, Alice S Serafin).

## Abstract

Nephronophthisis (NPH) is n rare recessive kidney disease caused by biallelic variants in more than 25 *NPHP* genes encoding proteins that localize to primary cilia. It is characterized by three different forms depending on the age of onset and kidney lesions: infantile (cystic), juvenile/late onset (fibrotic). To date, the pathways linking altered primary cilia function to progressive kidney scarring in NPH remain poorly defined and therapeutic options are lacking.

To address these questions, we generated two new mouse NPH models by inactivating *Nphp3* specifically in kidney tubules either during embryogenesis or in adult, recapitulating the infantile and juvenile forms of the disease, respectively. Embryonic inactivation produced a rapid and severe cystic phenotype with tubular dedifferentiation, progressive interstitial fibrosis, inflammation and kidney failure, while postnatal inactivation led to a slowly progressive tubulointerstitial nephropathy characterized by tubular atrophy, fibrosis and immune cell infiltration without cyst formation. Strikingly, cilia were preserved in the early stages of both models, indicating that ciliogenesis impairment is not a primary driver of NPH3 pathogenesis. Transcriptomic profiling of the juvenile model revealed that disease initiation is driven by mitochondrial dysfunction, innate immune activation and aberrant cell cycle progression, while *epithelial-to-mesenchymal transition* and *Wnt/β-catenin* remodelling emerges only at later stages of disease progression. Therapeutic intervention with the PGE1 (alprostadil) failed to rescue the cystic/infantile model but significantly attenuated fibrosis, inflammation and interstitial fibrosis in the fibrotic/juvenile model. The ability to recapitulate both disease forms through temporal modulation of gene inactivation suggests that primary cilia serve distinct, stage-specific functions in kidney tubular homeostasis, with different cellular processes being selectively vulnerable depending on the causative gene or variant. Collectively, these findings uncover early pathogenic mechanisms that may constitute tractable therapeutic targets for the treatment of nephronophthisis.

## INTRODUCTION

Nephronophthisis (NPH) is an autosomal recessive ciliopathy and tubulointerstitial nephropathy that stands as the leading monogenic cause of kidney failure in children and young adults (König et al. 2017; Tang et al. 2022). NPH presents either as an isolated kidney disease or in syndromic forms also affecting the liver, retina, and brain, among others. Based on the age at which kidney failure occurs, NPH is classified into three forms (Luo et Tao 2018; Wolf et Hildebrandt 2011). The juvenile form is the most prevalent, accounting for approximately 80% of cases, with kidney failure typically progressing around age 13. It is characterized by impaired urinary concentration, kidneys of normal or reduced size, tubular basement membrane abnormalities, tubular atrophy and/or dilatation, immune cell infiltration, and severe interstitial fibrosis. The adolescent/young adult form shares these histological features but manifests later in life. The infantile form is the earliest and most severe: symptoms appear within the first months of life, kidney failure occurs before age 5, and the kidneys are enlarged and studded with cortical cysts, already accompanied by interstitial fibrosis. Understanding why the same disease entity produces such divergent manifestations - from predominantly cystic kidneys in infants to fibro-inflammatory disease in juveniles - remains one of the central unresolved questions in the field. No disease-modifying therapy currently exists. Dialysis and kidney transplantation remain the only options for affected patients (Stokman, Saunier, et Benmerah 2021).

To date, biallelic variants in more than 25 genes encoding NPH proteins (NPHP) caused the disease, resulting in extensive genetic heterogeneity with overlapping clinical phenotypes. NPHP assemble into distinct complexes at the primary cilium, an organelle essential for the transduction of developmental and homeostatic signals. These complexes - including the NPHP1-4-8 module (Sang et al. 2011), which gates the ciliary transition zone, and the inversin compartment module (Bennett et al. 2020), which occupies the proximal ciliary shaft - are thought to maintain the highly specific molecular composition of cilia and to coordinate signalling within this compartment. Although the inversin compartment is required for correct cilia-dependent signalling in various tissues, notably the regulation of the Wnt/β-catenin pathway (Simons et al. 2005), its function in kidney tubular cells remained to be clarified. Disruption of these complexes can impair ciliogenesis and/or ciliary composition therefore altering signalling, including Hedgehog, polycystin-dependent flow sensing, Wnt, Hippo, and Akt. However, how perturbation of specific NPHP complexes translates into the distinct clinical and histological forms of NPH remains poorly understood.

NPHP3, encoding a key structural component of the inversin compartment, is the fourth most frequently affected gene in NPH, accounting for approximately 5% of diagnosed cases (Petzold et al. 2023). Within the inversin compartment, INVS (NPHP2), ANKS6 (NPHP16) and NEK8 (NPHP9) and NPHP3 assemble as fibrilloids at the proximal part of the axoneme functioning as a signalling scaffold (Hoff et al. 2013; Roig 2025). The disease spectrum associated with *NPHP3* variants spans from including isolated NPH - from infantile to adult forms - to severe foetal syndromic forms (Olbrich et al. 2003; Tory et al. 2009; Olinger et al. 2021; Petzold et al. 2023; Rajput et al. 2026). This breadth of phenotypic variation makes *NPHP3* a uniquely informative locus to dissect the mechanisms that determine NPH disease course. Yet, the cellular and molecular basis of this genotype-phenotype correlation remains entirely unexplained.

A major obstacle to progress in the field has been the absence of animal models that faithfully recapitulate the key histopathological hallmarks of human NPH, in particular the tubulointerstitial fibrosis and immune cell recruitment that dominates the juvenile form. Inactivation of *Nphp1*, the most common cause of juvenile NPH, does not lead to kidney failure in mice (Garcia, et al. 2022), and the same holds true for other NPHP orthologues (*Nphp4*). *Nphp7*/*Glis2* inactivation is a notable exception, consistently producing kidney fibrosis in mice (Kim et al. 2008; Lu et al. 2016a), but *NPHP7* variants account for fewer than 0.1% of human NPH cases (Petzold et al. 2023). For *NPHP3* specifically, existing rodent models occupy the two inadequate extremes of an incomplete spectrum. The pcy mouse carries a hypomorphic missense mutation in *Nphp3* (p.Ile614Ser) that preserves partial protein function; it develops a slowly progressive polycystic kidney disease but lacks the tubulointerstitial fibro-inflammation that defines juvenile NPH (Olbrich et al. 2003). At the opposite extreme, complete germline inactivation of *Nphp3* results in embryonic lethality in mice mirroring the most severe human *NPHP3* phenotypes but precluding any analysis of kidney dysfunction in adult animals (Lu et al. 2016). As a result, the impact of complete NPHP3 loss specifically in differentiated kidney tubular cells, and the influence of developmental timing on whether cystogenesis or fibro-inflammation ensues, has never been studied *in vivo*. This represents a fundamental gap.

Here, we report the generation and characterization of a conditional mouse model of *Nphp3* deficiency in which *Nphp3* was specifically inactivated in kidney tubular cells. We demonstrate that inactivation during kidney development produces a cystic phenotype recapitulating infantile NPH, whereas inactivation after kidney maturation results in progressive interstitial fibrosis and inflammation recapitulating juvenile NPH. This model provides, for the first time, an *in vivo* system capable of reproducing the full clinical spectrum of NPHP3-associated kidney disease and directly establishes that the timing of *Nphp3* loss in kidney tubules is a primary determinant of disease manifestation. Leveraging this model, we further evaluate the therapeutic efficacy of the prostaglandin analogue PGE1 in the context of fibro-inflammatory disease, placing its potential benefit in a clinically relevant pathological setting for the first time.

## RESULTS

### Embryonic *Nphp3* tubular inactivation induced severe cystic phenotype and kidney failure

We specifically inactivated *Nphp3* (*Nphp3*^ΔTub^) using KspCre that is expressed in the distal part of the tubules during embryonic day 10.5 (Shao, Somlo, et Igarashi 2002). *Nphp3* transcript was significantly decreased in kidneys from *Nphp3*^ΔTub^ mice as compared to control mice (**Supp Fig. 1A**). Macroscopically, the *Nphp3*^ΔTub^ kidneys increased in size over time reaching a 3-fold increase at postnatal day 21 (P21) compared to control mice, and ∼9-fold by P28 of age compared to control mice (**Fig. 1A, B** and **Supp Fig. 1B**). Consistent with advanced disease severity, *Nphp3*^ΔTub^ mice began to lose weight at P28 (**Fig. 1C**). Functional analyses revealed a progressive decline in kidney function: urinary osmolality was significantly impaired at P28 (**Fig. 1D**), while blood urea nitrogen (BUN) levels progressively increased starting at P21 in *Nphp3*^ΔTub^ compared to controls (**Fig. 1E**).

**Figure 1.**
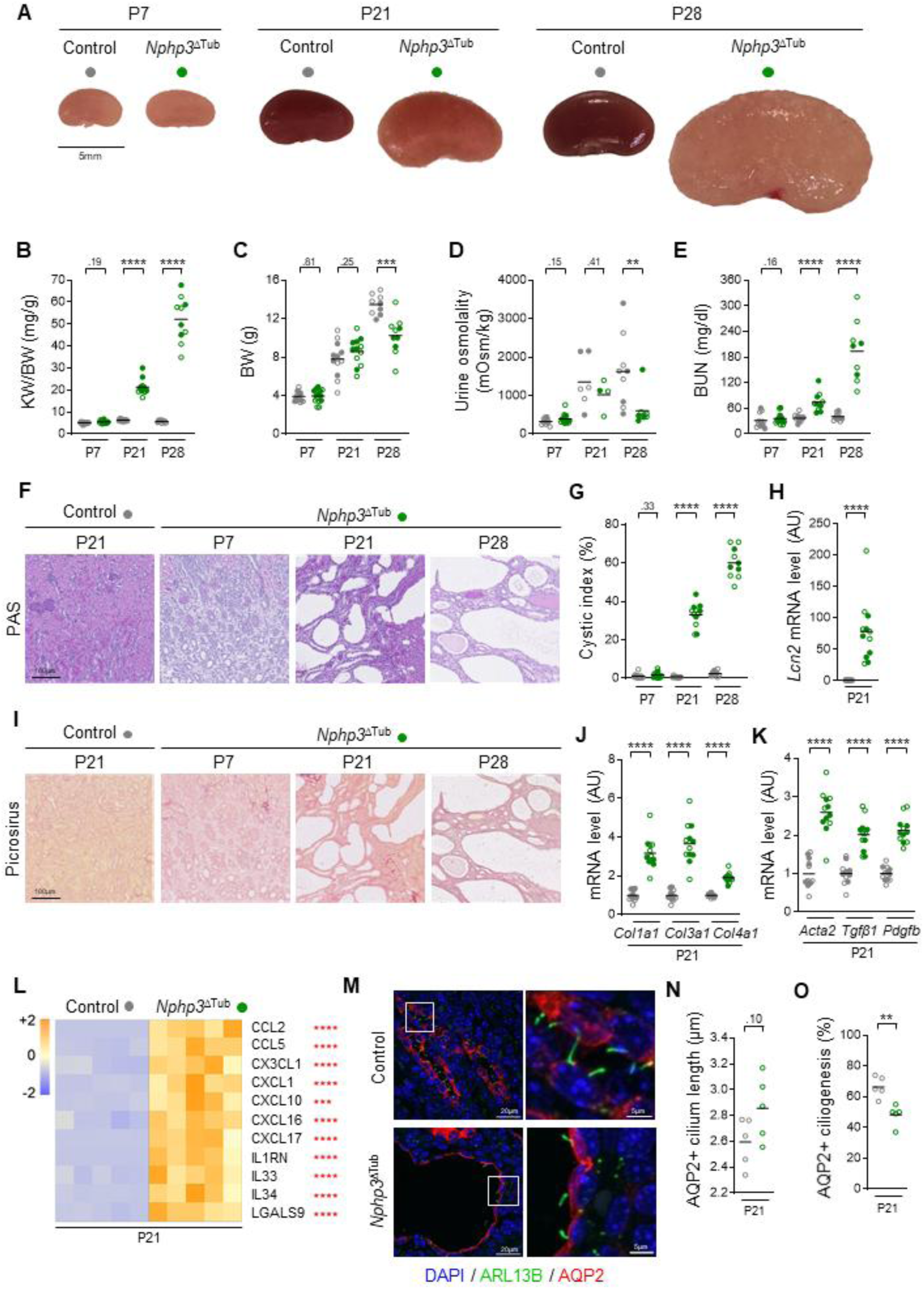
Embryonic *Nphp3* tubular inactivation induced severe cystic phenotype and kidney failure. **(A)** Representative kidney pictures from 7, 21 and 28 days-old control and *Nphp3*^ΔTub^ mice. Scale bar: 5mm. **(B-E)** Kidney weight (KW) to body weight (BW) ratio (B), body weight (C), urine osmolality (D) and blood urea nitrogen (BUN, E) from the same 6 mice groups. **(F, I)** Representative pictures of Periodic Acid Schiff (PAS, F) and Picrosirus (I) staining from 21 days-old control and 7, 21 and 28 days-old *Nphp3*^ΔTub^ mice. Scale bar: 100 µm. **(G)** Cystic index quantification from the same 6 mice groups. **(H, J-K)** *Lcn2* (H), *Col1a1, Col3a1* and *Col4a1* (J), *Acta2, Tgfβ1* and *Pdgfb* (K) mRNA expression in kidneys from the 21 days-old control and *Nphp3*^ΔTub^ mice. **(L)** Heatmap showing Z-scores computed on cytokines mRNA expression in kidneys from the 21 days-old control and *Nphp3*^ΔTub^ mice **(also in Supplementary** Fig. 1N**-X)**. **(M-O)** Representative images (M) and quantification of primary cilia (ARL13B+, green) length (N) and density (O) in collecting ducts (CD, AQP2+, red) of kidneys from 21 days-old control and *Nphp3*^ΔTub^ mice. Scale bar: 20μm (left panels) and 5µm (right panel). **(B-E, G-H, J-L, N-O)** Each dot represents one individual mouse. Empty circle for males, filled circles for females. Bars indicates mean. Mann-Whitney test: ** *P* < 0.01, \*\*\**P* < 0.001, \*\*\*\**P* < 0.0001. AU: arbitrary unit.

Histology revealed fluid-filled cysts (**Fig. 1F** and **Supp Fig. 1C**) with slight tubular dilation at P7 and a progressively increasing cyst burden over time (**Fig. 1G**). This cystic expansion was paralleled by a marked upregulation of the tubular injury marker Lipocalin2 (*Lcn2*) mRNA level (**Fig. 1H**) and by a broad disruption of tubular differentiation markers (**Supp Fig. 1D-H**). Progressive interstitial fibrosis was also observed with deposition of collagen fibers and increased expression of pro-fibrotic markers (collagens, *Acta*2, *Tgfb1, Pdgfb*) in *Nphp3*^ΔTub^ (**Fig. 1I-K**). From P21 onward, *Nphp3*^ΔTub^ kidneys also presented with enhanced NPH-associated pro-inflammatory cytokines (**Fig. 1L** and **Supp. Fig. 1I-S**) (Quatredeniers et al. 2022).

As NPHP3 is a ciliary protein, we investigated whether its invalidation influenced ciliogenesis. Cilia length remained unchanged in *Nphp3*^ΔTub^ kidneys as compared to control ones (**Fig. 1M, N**). However, once cysts expanded, at P21, we observed a significant ciliogenesis defect in AQP2-positive tubular cells of *Nphp3*^ΔTub^ kidneys as compared to controls (**Fig. 1O**).

In conclusion, loss of *Nphp3* tubular epithelial cells during embryogenesis results in fluid-filled cysts, tubular dedifferentiation, and chronic inflammation with progressive scarring leading to kidney failure, reminiscent of the infantile NPH in humans.

### Adult *Nphp3* tubular inactivation induced tubular atrophy, interstitial fibrosis, and inflammation

To induce a postnatal invalidation of *Nphp3*, we took advantage of the tamoxifen-inducible kidney tubule-specific Cre recombinase (KspCad-CreER^T2^) (Lantinga-van Leeuwen et al. 2006). Tamoxifen was administered at one month of age to conditionally inactivate *Nphp3* in kidney tubules (i*Nphp3*^ΔTub^). *Nphp3* transcript was significantly decreased in kidneys from i*Nphp3*^ΔTub^ mice as compared to controls already 2 months after invalidation (3-month-old mice) (**Supp Fig. 2A**). No differences in behavior or body weight were observed between i*Nphp3*^ΔTub^ and control animals (**Fig. 2A**). Macroscopically, the i*Nphp3*^ΔTub^ kidneys looked normal in shape; however, over time a significant increase in kidney weight to body weight ratio was observed at 10 months of age (**Fig. 2B, C**). Histology revealed a slowly progressive kidney phenotype with tubular atrophy, tubular dilatation and immune cell infiltration (**Fig. 2D**), associated with increased *Lcn2* injury marker expression and tubular dedifferentiation in i*Nphp3*^ΔTub^ kidneys (**Fig. 2E** and **Supp. Fig. 2B-F**). Transmission electron microscopy revealed thickened tubular basement membranes in 10-month-old i*Nphp3*^ΔTub^ as compared to controls (**Fig. 2F**). Interstitial fibrosis emerged as a prominent feature: collagen deposits and mRNA expression levels were significantly increased in i*Nphp3*^ΔTub^ kidneys (**Fig. 2G-H**) and were associated with enhanced expression of pro-fibrotic markers (*Acta2*, *Tgfb1*, *Pdgb*; **Fig. 2G-J** and **Supp Fig. 2G-L**). Consistent with the inflammatory signature described in tubular cells derived from NPH patients (Quatredeniers et al. 2022), i*Nphp3*^ΔTub^ mice exhibited elevated expression of *Cd3*, *Cd4*, *Adgre1* and NPH-associated pro-inflammatory cytokines (**Fig. 2K** and **Supp Fig. 3E-O**).

**Figure 2.**
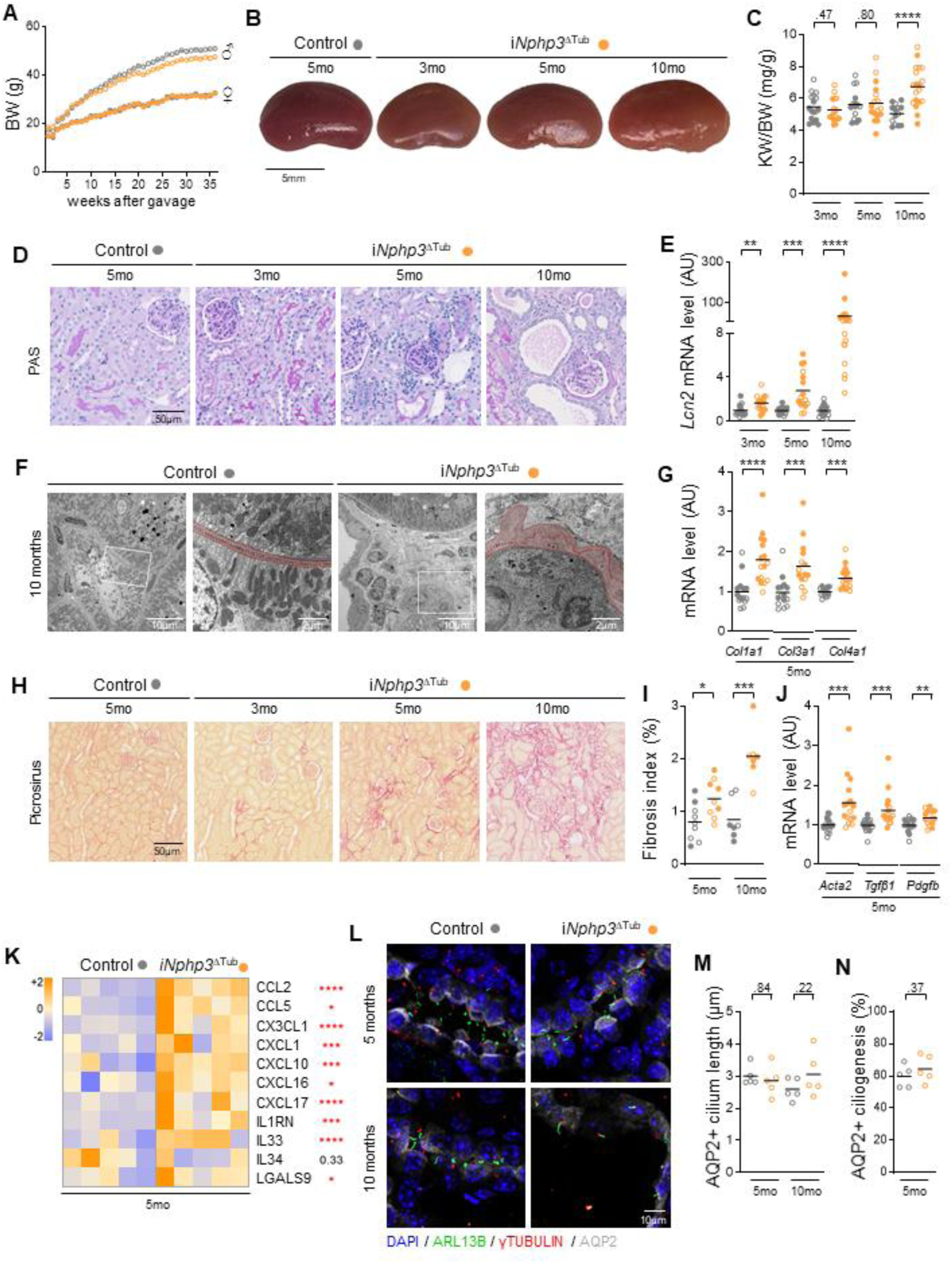
Adult *Nphp3* tubular inactivation induced tubular atrophy, interstitial fibrosis, and inflammation. **(A)** Body weight (BW) follow-up curve from 1 to 36 weeks after tamoxifen gavage from 10 months-old control and i*Nphp3*^ΔTub^ mice. **(B)** Representative kidney pictures from 5 months-old control and 3, 5 and 10 months-old i*Nphp3*^ΔTub^ mice. Scale bar: 5mm. **(C)** Kidney weight (KW) to body weight (BW) ratio from 3, 5 and 10 months-old control and i*Nphp3*^ΔTub^ mice. **(D)** Representative pictures of Periodic Acid Schiff (PAS) staining from 5 months-old control and 3, 5 and 10 months-old i*Nphp3*^ΔTub^ mice. Scale bar: 50µm. **(E)** *Lcn2* mRNA expression in kidneys from the 6 same mice groups. **(F)** Representative pictures of transmission electron microscopy from 10 months-old control and i*Nphp3*^ΔTub^ mice with the basement membrane highlight in red. Scale bar: 10µm and 2µm (zoomed-in panels). **(G)** *Col1a1*, *Col3a1* and *Col4a1* mRNA expression in kidneys from the 5 months-old control and i*Nphp3*^ΔTub^ mice. **(H-I)** Representative pictures of Picrosirus staining (H) and fibrosis quantification (I) from 5 months-old control and 3, 5 and 10 months-old i*Nphp3*^ΔTub^ mice. Scale bar: 50µm. **(J)** *Acta2, Tgfβ1* and *Pdgfb* mRNA expression in kidneys from the 5 months-old control and i*Nphp3*^ΔTub^ mice. **(K)** Heatmap showing Z-scores computed on cytokines mRNA expression in kidneys from the 5 months-old control and i*Nphp3*^ΔTub^ mice **(also in Supplementary** Fig. 3E**-O)**. Mann-Whitney test: * *P* < 0.05, \*\*\**P* < 0.001, \*\*\*\**P* < 0.0001. **(L-N)** Representative images and quantification of primary cilia (AcTUB+, green) length (M) and density (only at 5 months, N) in collecting ducts (CD, AQP2+, grey) of kidneys from 5 and 10 months-old control and i*Nphp3*^ΔTub^ mice. Scale bar: 10μm.

Cilia were also assessed: neither ciliogenesis nor cilia and inversin compartment lengths were altered in i*Nphp3*^ΔTub^ kidneys (**Fig. 2L**, **Supp Fig. 3P**).

Altogether, these results show that loss of *Nphp3* in tubular cells after the end of kidney maturation results in tubular atrophy, dilation, and dedifferentiation as well as tubular basement thickening. It also leads to chronic inflammation with interstitial fibrosis, reminiscent of the juvenile NPH.

### Adult *Nphp3* tubular inactivation induced decreased mitochondrial biogenesis and enhanced inflammation and cell cycle

To explore the molecular mechanisms triggered by the loss of *Nphp3* in the juvenile NPH mouse model, bulk RNA-seq was conducted on kidneys from i*Nphp3*^ΔTub^ and control mice at 3 months, an early stage in the onset of the disease, and at 5 months, a stage at which kidney lesions are already well established (**Fig. 2**). *Nphp3* inactivation induced 40 differential expressed genes (DEGs) at 3 months and 106 at 5 months, of which 6 were commonly deregulated (**Fig. 3A, B**). From these 6 common DEGs, *Nphp3* was down-regulated as expected, while *Havcr1*, *Ccl2*, *Cd44*, *Serpina10* and *Aoc1* were upregulated in i*Nphp3*^ΔTub^ kidneys (**Fig. 3A**). Even though the number of DEGs were not so high, gene set enrichment analysis (GSEA) of our transcriptomic data revealed that, at disease onset, NPHP3 loss primarily affected three major functional modules : (i) mitochondrial dysfunction associated with a metabolic shift toward aerobic glycolysis at the latest stage (*Oxidative Phosphorylation*, *Glycolysis*), (ii) activation of inflammatory pathways (*Complement*, *TNFa signalling*, *Interferon signalling*, *IL6/JAK/STAT3 signalling*, *inflammatory response*) and (iii) dysregulated proliferative activity (*E2F targets*, *G2/M checkpoint*, *Mitotic spindle*) (**Fig. 3C**). Interestingly, the *Epithelial Mesenchymal Transition* hallmark emerged predominantly at the latest disease stage (5 months), as well as the *Apical Junction* and *Wnt beta catenin signaling* hallmarks, suggesting that epithelial remodeling and pro-fibrotic transcriptional programs are features of advanced disease progression rather than early pathogenic events (**Fig. 3C, D**, upper). GSEA also confirmed that NPHP3 loss did not induced dysregulation of ciliary genes specific to the kidney (*Kidney ciliome*, **Fig. 3D**, lower and **Supp Table 2**).

**Figure 3.**
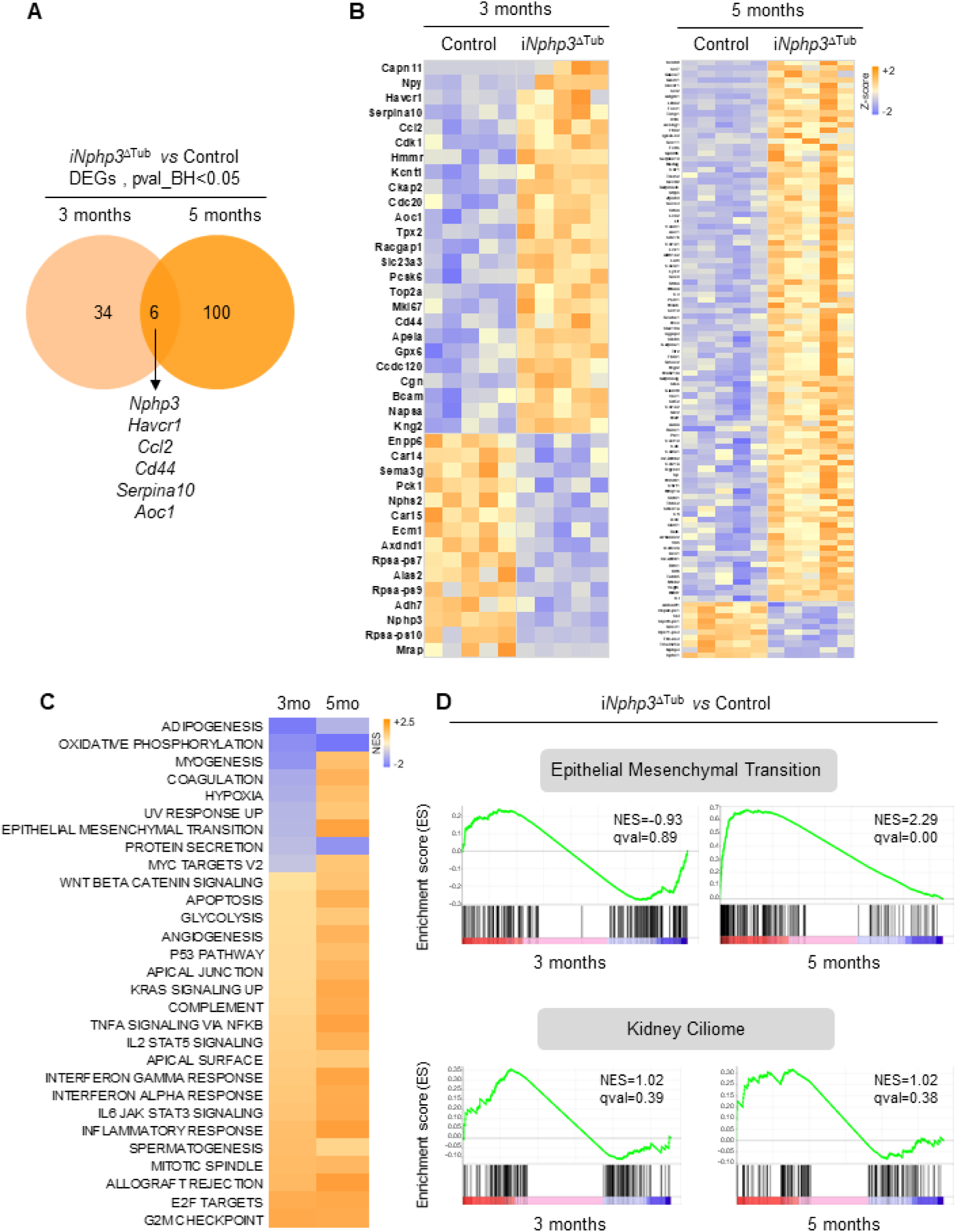
Adult *Nphp3* tubular inactivation induced decreased mitochondrial biogenesis and enhanced inflammation and cell cycle. (A-B) Venn diagram (A) representing the differentially expressed genes (DEGs) in i*Nphp3*^ΔTub^ mice compare to control at 3 months (left circle), 5 months (right circle) and the 6 common DEGs (intersection). These 40 (3 months) and 106 DEGs (5 months) are represented on the heatmaps showing the computed Z-scores of the normalized raw counts (B). **(C)** Heatmaps computed on the normalized enriched score (NES)-ranked gene sets (MSigDB hallmarks, GSEA) in 3 and 5 months i*Nphp3*^ΔTub^ mice compared to controls. Only pathways with FDR<0.05 at one time point are represented. **(D)** Enrichment plot from GSEA analysis of the *Epithelial Mesenchymal Transition* hallmark genes and the designed *Kidney Ciliome* (comprising 151 genes) comparing the i*Nphp3*^ΔTub^ mice to control at 3 months (left panel) and 5 months (right panel).

In conclusion, transcriptomic analysis of the NPHP3 juvenile model revealed that initiation of the disease involved metabolic stress, inflammation and cell-cycle progression, while during progression of the disease, tubular cells lose their epithelial characteristics and acquired a “mesenchymal-like” feature.

### Alprostadil treatment ameliorates kidney tubulointerstitial lesions in adult NPHP3 loss model but not in the embryonic one

We took advantage of these two mouse models recapitulating infantile NPH with cystic phenotype and juvenile NPH with fibro-inflammation phenotype, to test the therapeutic effect of the PGE2 analog PGE1, alprostadil (ALPRO), which showed efficacy in reversing ciliogenesis defects and tubular dilations associated with NPHP1 loss in patient-derived kidney tubular cells and a KO mouse model (Garcia, et al. 2022).

For the infantile model (embryonic *Nphp3* inactivation), ALPRO was administrated intraperitoneally daily (80μg/kg mouse) from P10 to P21 to *Nphp3*^ΔTub^ and control mice without any impact on growth or survival. Macroscopic and histologic analysis of the kidneys revealed no beneficial improvement of cystic index after ALPRO treatment (**Supp Fig. 4A-C**). Urine concentration and kidney function were still impaired in *Nphp3*^ΔTub^ mice treated with ALPRO in comparison to DMSO-treated *Nphp3*^ΔTub^ mice (**Supp Fig. 4D-E**). In line with these results, no improvement of collagen deposits or fibrotic markers levels or immune markers levels was measured in ALPRO-treated *Nphp3*^ΔTub^ mice (**Supp Fig. 4F-K**).

For the juvenile model (adult *Nphp3* inactivation), ALPRO was administrated intraperitoneally (80ug/kg mouse) daily from 6 week to 5 months to i*Nphp3*^ΔTub^ and control mice without any impact on growth or survival. Histology and qPCR revealed a slight but significant improvement of the progression of the disease, with less fibrosis (*Col1a1*, *Acta2*), inflammation (*Cd3*, *Adgre1*) and inflammatory cytokines (*Ccl5*, *Cxcl10*) in ALPRO-treated i*Nphp3*^ΔTub^ mice (**Fig. 4A-H**). GSEA of bulk-RNAseq from DMSO- or ALPRO-treated i*Nphp3*^ΔTub^ and control kidneys revealed that ALPRO treatment efficiently reversed the pathways involved in the initiation (Cell cycle and Inflammation) and the progression (Epithelial Mesenchymal transition) of the disease (**Fig. 4I**).

**Figure 4.**
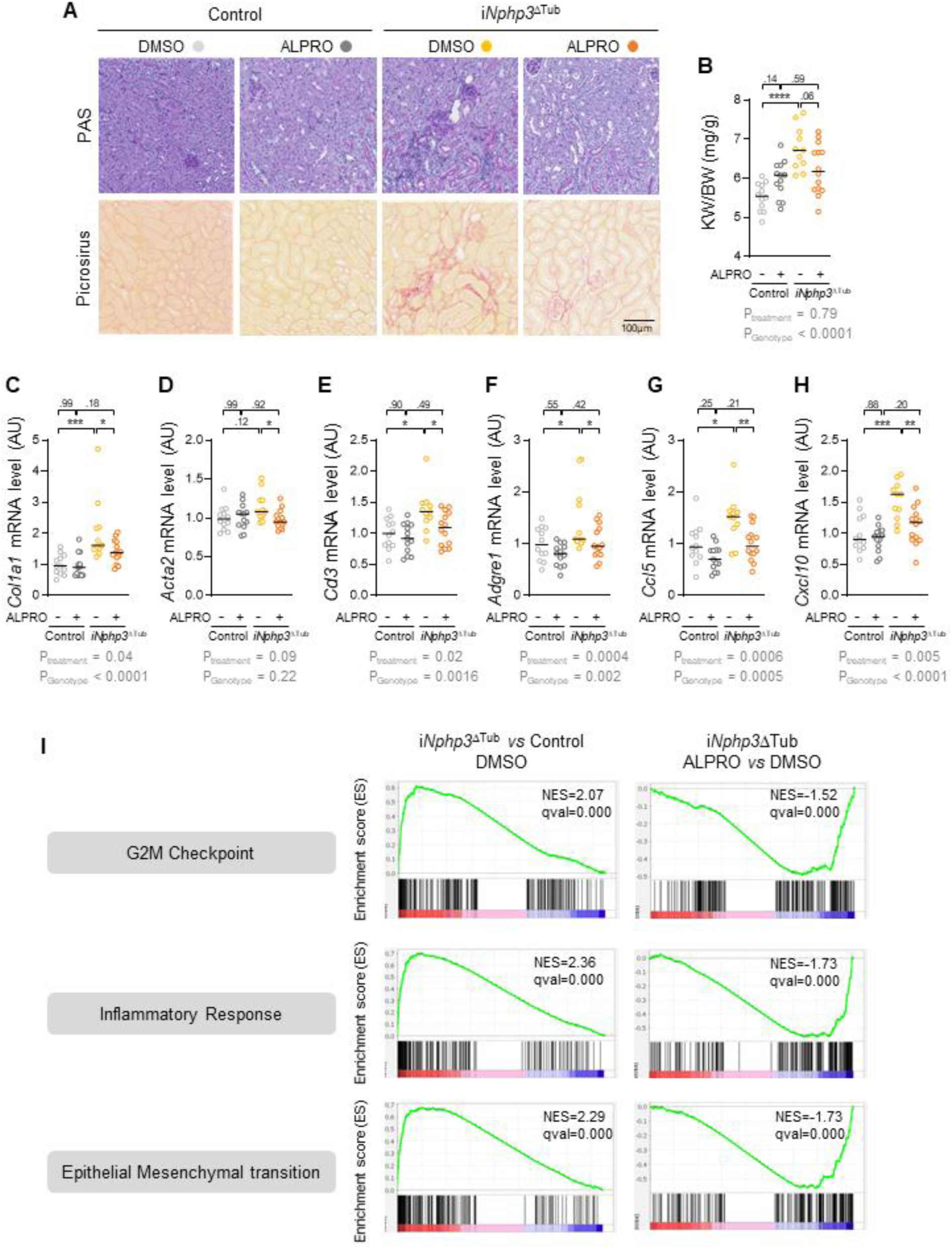
Alprostadil treatment ameliorates kidney tubulointerstitial lesions in *Nphp3* juvenile mouse model. **(A)** Representative pictures of Periodic Acid Schiff (PAS) and Picrosirus staining from 5 months-old control and i*Nphp3*^ΔTub^ mice treated with DMSO or Alprostadil (ALPRO). Scale bar: 100 µm. **(B)** Kidney weight (KW) to body weight (BW) ratio from the same 4 mice groups. **(C-H)** *Col1a1* (C), *Acta2* (D), *Cd3* (E), *Adgre1* (F), *Ccl5* (G) and *Cxcl10* (H) mRNA expression in kidneys from the same 4 mice groups. Each dot represents one individual mouse. Empty circle for males. Bars indicates mean. Two-ways anova test: \**P* < 0.05, ** *P* < 0.01, \*\*\**P* < 0.001, \*\*\*\**P* < 0.0001. *P_treatment_* corresponding to P-value of the treatment affect and *P_Genotype_* corresponding to the P-value of the genotype affect. AU: arbitrary unit. **(I)** Enrichment plot from GSEA analysis of the “G2M Checkpoint”, “Inflammatory Response” and “Epithelial Mesenchymal Transition” hallmarks comparing the i*Nphp3*^ΔTub^ mice to control mice treated with DMSO at 5 months (left panel) and i*Nphp3*^ΔTub^ mice treated with ALPRO to i*Nphp3*^ΔTub^ mice treated with DMSO (right panel).

## DISCUSSION

In this study, we generated two complementary mouse models of nephronophthisis by conditionally inactivating *Nphp3* either during embryogenesis or in adult kidney tubules, recapitulating the infantile and juvenile clinical forms of the disease, respectively. Together, these models provide new insights into the temporal pathogenesis of NPH and reveal a differential response to therapeutic intervention with the PGE1 analog alprostadil.

The embryonic tubular inactivation of *Nphp3* (*Nphp3*^ΔTub^) produced a rapid and severe cystic phenotype consistent with infantile NPH. Kidneys enlarged dramatically within the first weeks of life, accompanied by progressive loss of concentrating ability, rising BUN levels, and early kidney failure. These features closely mirror the aggressive clinical course observed in patients carrying biallelic *NPHP3* mutations presenting in infancy (Tory et al. 2009; Petzold et al. 2023). Histologically, tubular dilations emerged as early as one week of age and expanded into fluid-filled cysts, parallel with tubular dedifferentiation, injury marker upregulation (Lcn2), interstitial fibrosis, and a prominent pro-inflammatory cytokine signature. The early onset and rapid cystic expansion in this model likely reflect the critical role of NPHP3 during the period of active nephron maturation, when tubular cells are particularly sensitive to disruption of ciliary signalling and planar cell polarity pathways that govern tubulogenesis (Fischer et al. 2006; Piontek et al. 2007; Patel et al. 2008). Notably, while cilia length was preserved at early stages, a secondary ciliogenesis defect emerged in AQP2-positive collecting duct cells once cysts had expanded, suggesting that ciliogenesis impairment in this context is a consequence rather than a primary driver of cyst formation - a distinction with important mechanistic implications.

In contrast, postnatal tubular inactivation of *Nphp3* (i*Nphp3*^ΔTub^) produced a slowly progressive phenotype reminiscent of juvenile NPH, the most common form of the disease in humans. The absence of cyst formation in this model and the gradual accumulation of tubular atrophy, basement membrane thickening, interstitial fibrosis and immune cell infiltration over months is consistent with the clinical presentation of juvenile NPH, which typically manifests with interstitial nephritis and fibrosis rather than cystic disease (Petzold et al. 2023).

Also, the two orthologous *Nphp3* models described here induced bi-allelic tubular loss of function of *Nphp3*, which in humans leads to severely enlarged kidneys, timely inactivation uncovered distinct functions of NPHP3. These observations reinforce the emerging concept that during development, NPHP3 is necessary for correct tubulogenesis - as a component of the primary cilia machinery - while once kidney development is complete, NPHP3 is required for maintaining tubular epithelial integrity. These findings also suggest that, although the mature tubular epithelium is no longer prone to cystic transformation, it remains dependent on NPHP3 function for the maintenance of tubular integrity and homeostasis. Together, these models represent unique tools to dissect NPHP3 function in tubular homeostasis.

Transcriptomic profiling of the juvenile model at two disease stages provided mechanistic insight into the sequence of pathological events following *Nphp3* loss. At the earliest timepoint (3 months), gene set enrichment analysis identified mitochondrial dysfunction, with a subsequent metabolic shift toward aerobic glycolysis, alongside activation of inflammatory pathways and enhanced cell cycle progression as the dominant dysregulated modules. This metabolic reprogramming is reminiscent of the Warburg-like shift described in other tubular injury settings and in polycystic kidney disease (Rowe et al. 2013), and may reflect an adaptive but ultimately maladaptive response to energetic stress caused by NPHP3 deficiency. The early and prominent inflammatory signature, encompassing complement activation, TNFα, interferon, and IL-6/JAK/STAT3 signalling, is consistent with the pro-inflammatory gene expression profile previously described in tubular cells derived from NPH patients (Quatredeniers et al. 2022), and positions innate immune activation as an early and potentially primary event in NPHP3 pathogenesis rather than a secondary consequence of fibrosis. The upregulation of cell cycle progression genes at this stage may reflect a tubular regenerative response to injury, though aberrant proliferation in post-mitotic tubular epithelial cells can itself promote dedifferentiation and disease progression (Lee, Gusella, et He 2021; Zhang et al. 2021).

Confirming human juvenile NPH biopsies, the *Epithelial-to-Mesenchymal Transition* (EMT) hallmark emerged predominantly at the later disease stage (5 months) in i*Nphp3*^ΔTub^ mice, together with dysregulation of Wnt/β-catenin and apical junction pathways. This temporal ordering suggests that epithelial remodelling and the acquisition of a mesenchymal-like transcriptional program are features of disease progression rather than initiating events, and likely reflect the downstream consequences of sustained metabolic stress and chronic inflammation. A similar temporal dissociation between early metabolic/inflammatory injury and late fibrotic remodelling has been proposed in other chronic kidney disease models (Venkatachalam et al. 2015; Xu et al. 2022; Miguel, Shaw, et Kramann 2025), and our data provide direct transcriptomic evidence for this hierarchy in the context of NPH3. The differential response of the two models to alprostadil treatment is mechanistically informative. Alprostadil, a PGE1 analog previously shown to rescue ciliogenesis defects and tubular dilation in *Nphp1*-deficient mice and patient-derived tubular cells (Garcia, et al. 2022), failed to improve any parameter of disease in the embryonic *Nphp3*^ΔTub^ model. This lack of efficacy may reflect the severity and rapidity of cyst formation in the infantile model, which may overwhelm any pharmacological intervention administered after cysts have already initiated, or may indicate that the pathogenic mechanisms operating during embryonic tubulogenesis are fundamentally distinct from those targeted by alprostadil. In contrast, alprostadil produced a significant, albeit modest, amelioration of the juvenile phenotype, reducing fibrosis, immune cell infiltration and inflammatory cytokine expression in *Nphp3*^ΔTub^ mice. Importantly, transcriptomic analysis confirmed that alprostadil reversed the gene expression programs associated with both early disease initiation (inflammation and cell cycle) and later disease progression (EMT), suggesting that its therapeutic benefit extends beyond ciliogenesis rescue and actin remodelling and may involve broader anti-inflammatory and anti-fibrotic mechanisms. The precise molecular targets through which alprostadil exerts these effects in the context of NPHP3 loss remain to be fully elucidated.

Several limitations of this study should be noted. The bulk RNA-sequencing approach, while providing pathway-level resolution, does not capture cell-type-specific transcriptional changes within the tubular and interstitial compartments. Single-cell or single-nucleus RNA-sequencing at multiple disease stages would provide higher resolution of the cellular dynamics underlying NPH progression. Additionally, while alprostadil showed efficacy in the juvenile model, the effect size was moderate, suggesting that combination approaches targeting multiple disease pathways - for instance, pairing alprostadil with anti-inflammatory or anti-fibrotic agents - may be required for more substantial therapeutic benefit.

In conclusion, our findings establish that NPHP3 plays distinct roles in the developing versus the mature kidney, with its loss producing clinically and molecularly distinct disease phenotypes depending on the timing of inactivation. We identify metabolic stress, innate immune activation, and cell cycle dysregulation as early and potentially targetable events in NPHP3 pathogenesis, and demonstrate that pharmacological intervention with alprostadil can partially reverse these pathological programs in the juvenile disease context. These results have translational implications for the stratification and treatment of NPH patients based on disease onset and molecular disease stage.

## METHODS

### Mice

Mice were housed in a specific pathogen-free facility, fed ad libitum and housed at constant ambient temperature in a 12-hour day/night cycle. Breeding and genotyping were done according to standard procedures.

To generate *Nphp3*^flox/flox^ mice (mixed C57BL/6N x DBA2/J background), we used CRISPR/Cas9 technology to insert LoxP sites in the introns before exon 4 and after exon 5, respectively (Rajput, et al. 2026). In this way, Cre-mediated recombination induced the deletion of exons 4 and 5 and created a frameshift leading to the appearance of a premature stop codon causing a loss of 83% of NPHP3 protein sequence. CRISPR direct software (http://crispr.dbcls.jp/) was used for designing the guides RNA. Two gRNAs were designed to target DNA sequences in introns 3 and 4. Two long single-strand oligonucleotides containing homology arms and LoxP site were synthesized as ultramer and used as repair template for HDR. Transgenesis steps related to pronuclear micro-injection of fertilized oocytes and transfer of injected zygotes into oviduct of pseudo-pregnant female mice were performed as above. Identification of the positive founders with insertion of both LoxP sites was done by PCR genotyping and confirmed by Sanger sequencing. The sequence of genotyping primers for the upstream LoxP site was as follows: sense primer 5’-GCT AAA GTG CTC TGC CAA A-3’, and antisense primer 5’-GGA TCA TGC TGC CTA TGG AAT-3’. The sequence of genotyping primers for the downstream LoxP site was as follows: sense primer 5’-GAT GAA ACA CAG CCC GAA ATG-3’, and antisense primer 5’-GCC TGC ATG TGT ACT CTG T-3’. The annealing conditions were 52°C for 1 minutes 30 seconds.

For developmental *Nphp3* inactivation, *Nphp3*^flox/flox^ mice were backcrossed for 2 generations with Ksp-Cre (B6.Cg-Tg(Cdh16-cre)91lgr/J; mixed C57Bl/6N; C57Bl/6J background) (Shao et al. 2002) and then intercrossed to generate distal tubule-specific *Nphp3* knockout (further referred to as *Nphp3*^ΔTub^ mice). Experiments were conducted on both females and males.

For *Nphp3* inactivation at an adult stage, *Nphp3*^flox/flox^ mice were backcrossed for 2 generations with KspCad-CreER^T2^ (Tg(Cdh16-cre/ERT2)F427Djmp; C57BL/6J background) (Lantinga-van Leeuwen et al. 2006) and then intercrossed to generate Tamoxifen-inducible kidney-specific *Nphp3* knockout (further referred to as i*Nphp3*^ΔTub^ mice). One-month-old mice received tamoxifen (225µg/g, dissolved in 85% sunflower seed oil and 15% ethanol) for 5 consecutive days with a feeding needle. Experiments were conducted on both females and males.

Alprostadil treatment was performed on both *Nphp3*^ΔTub^ (males and females) and i*Nphp3*^ΔTub^ (only males) mice. Mice received intraperitoneal injection of Alprostadil (80µg/kg) or vehicle (DMSO 0.04%) daily from P10 to P21 for *Nphp3*^ΔTub^ mice and from 6 weeks to 5 months for i*Nphp3*^ΔTub^ animals.

Littermates lacking Cre recombinase transgene were used as controls.

### Urine and Plasma Analyses

For *Nphp3*^ΔTub^ mice, spot urine samples were collected. For i*Nphp3*^ΔTub^ mice, at 6 months-old and 10 months-old, 24-hour urine samples were obtained from mice housed in metabolic cages. Urine excretion was measured. Urine osmolality was measured with a freezing point depression osmometer (Micro-Osmometer from Knauer).

Retro-orbital blood was collected from anesthetized mice. Plasma blood urea nitrogen (BUN) was measured using a Konelab 20i Analyzer (Thermo Scientific).

### Morphological Analysis

Kidneys were fixed in 4% paraformaldehyde, embedded in paraffin, and 4µm sections were stained with periodic acid-Schiff (PAS) or Picrosirius Red. Stained full size images were recorded using a whole slide scanner Nanozoomer 2.0 (Hamamatsu) equipped with a 20x/0.75 NA objective coupled to NDPview software (Hamamatsu). For *Nphp3*^ΔTub^, cystic index was measured using FIJI software from full sized kidney images of PAS staining and visualized at the percentage of all the empty circular area to the total kidney section area.

For fibrosis quantification, PicroSirius Red stained area was measured with FIJI software from full sized kidney images and visualized as the percentage of stained surface to total kidney section area.

### Transmission electron microscopy

Kidneys from 10-month-old i*Nphp3*^ΔTub^ animals were cut in small pieces (1mm^3^) and fixed with 2.5% glutaraldehyde in 0.1 M sodium cacodylate buffer prepared at pH 7.4 for 24 hours at 4°C. After rinsing with 0.2 M sodium cacodylate buffer, kidney samples were maintained in the same buffer at 4°C. Subsequently, they were post-fixed in 1% OsO4, dehydrated, and embedded in epoxy resin blocks. Blocks were then cut with an UC7 ultramicrotome. Finally, 70 nm-thick sections were recovered on copper and visualized with Hitachi HT7700 electron microscope. Pictures (2048x2048 pixels) of PTs, DCTs and CDs were imaged with an AMT41B and processed using ImageJ2. TEM acquisition was performed by the Imaging Facility of Hôpital Tenon (INSERM UMRS_S1155).

### Quantitative RT-PCR

Total RNAs were obtained from mouse kidneys using RNeasy Mini Kit (Qiagen) and reverse transcribed using High-Capacity cDNA Reverse Transcription Kit (Applied Biosystems) according to the manufacturer’s protocol. Quantitative PCR were performed with iTaq™ Universal SYBR® Green Supermix (Bio-Rad) on a CFX384 C1000 Touch (Bio-Rad). *Ppia*, *Rpl13* and *Sdha* were used as normalization controls (Vandesompele et al. 2002). Each biological replicate was measured in technical duplicates. The primers used for qRT-PCR are listed in **Supplementary Table 1**. Heatmaps displaying Z-scores computed on the expression levels of the identified cytokines, measured by qPCR, were generated using excel.

### Bulk RNA-seq

#### Samples preparation and sequencing

Total RNAs were extracted and purified from mouse kidneys using RNeasy Mini Kit (Qiagen), RNA quality was verified by capillary electrophoresis using Fragment AnalyzerTM. Bulk mRNA-Seq libraries were prepared by the Imagine Genomic Core Facility (Paris, France) starting from 1µg of total RNA, using the NEBNext® Low Input RNA Library Prep Kit according to manufacturer’s guidelines. This kit generates mRNA-Seq libraries from the PolyA+fraction of the total RNA by template switch reverse transcription. The cDNA was amplified twice and the final mRNAseq libraries are not ‘oriented’ or ‘stranded’ (meaning that the information about the transcribed DNA strand is not preserved during the library preparation). The equimolar pool of libraries, assessed by Q-PCR KAPA Library Quantification kit (Roche) and with a run test using the iSeq100 (Illumina) was sequenced on a NovaSeq6000 (Illumina, S2 FlowCells, paired-end 100+100 bases, ∼50 millions of reads/clusters produced per library). After the demultiplexing and before the mapping, the reads were trimmed to remove the adaptor sequences from the first amplification. *Differentially expressed genes analysis*: FASTQ files were mapped to the GRCCm38 primary mouse genome assembly using STAR and counted by featureCounts from the Subread R package. Normalization was performed with R package DESeq2 v1.24.0 (Love, Huber, et Anders 2014). For differential gene expression (DEGs) analysis, a Benjamini-Hochberg correction was applied with threshold for significance set as adjusted P-values < 0.05.

#### Gene set enrichment analysis (GSEA)

GSEA was performed on normalized raw counts according to the software manual (https://www.gsea-msigdb.org/) (Reimand et al. 2019). The following gene-set collections were included in the analysis: mouse hallmark (H), canonical pathways (C2:CP) and ontology (C5:BP, CC, MF). All parameters for GSEA were left as default. Gene sets with FDR<0.05 were considered significant. A kidney cilia gene-set (“kidney ciliome”) was designed as the genes expressed in the mouse kidney (Jensen et al. 2019) and present in Cilia Carta (Van Dam et al. 2019), SYSCILIA gold standard (Vasquez, van Dam, et Wheway 2021) and CilioGenics (151 genes, **Supplementary Table 2**).

### Immunofluorescence (IF)

7µm sections of paraffin-embedded kidneys were used for IF. Sections were first submitted to paraffin removal and antigen retrieval treatment (10 mM Tris pH9 – 1 mM EDTA – 0.05% Tween-20). For IHC, ANSK6 staining by IF, we performed 3% H_2_O_2_ (Sigma, #95321) and avidin/biotin blocking (Vector, #SP-2001) steps. Sections were then blocked with TBST – 10% FBS for 20 minutes and incubated overnight at 4 °C with the indicated primary antibodies. The following day, 1h room temperature incubation of the secondary antibody and nuclear staining was performed. Antibodies used are listed in **Supplementary Table 3**.

### Data availability

The transcriptomic data supporting the findings of this study will be openly available in the public domain. All the mice and cell models used in this study can be obtained upon request to the corresponding author, with the agreement of the scientist who generated the initial transgenic line.

### Statistical Analysis

Data were expressed as means. When only two groups were compared, differences between groups were evaluated using Student’s Mann-Whitney test. When testing more than 2 groups, two-way ANOVA followed by Tukey’s multiple comparisons test was used. The statistical analysis was performed using GraphPad Prism V10 software. All image analyses and mouse phenotypic analyses were performed in a blinded fashion.

### Study Approval

All animal experiments were conducted according to the guidelines of the National Institutes of Health Guide for the Care and Use of Laboratory Animals, as well as the French laws for animal welfare, and were approved by regional authorities (Ministère de l’Enseignement, de la Recherche et de l’Innovation # 32143, 39771 and 54621).

### Transparency

Artificial intelligence (https://claude.ai/) was used for editing clarity and grammar and language polishing.

## ACKNOWLEDGEMENTS

We are grateful to Dorien J.M. Peters (Department of Human Genetics, Leiden University Medical Center, Leiden, the Netherlands) for sharing Tamoxifen-inducible kidney-specific mice (KspCad-CreER^T2^) (Lantinga-van Leeuwen et al. 2006).

We thank the members of the LEAT, histology, genomics, bioinformatics and cell imaging facilities (S.F.R Necker INSERM US24, CNRS UAR 3633 Paris, France) and of the mouse renal physiology facility (Cordeliers Research Center, Paris, France) for technical assistance.

## SUPPLEMENTARY FIGURES

**Supplementary Figure 1.**
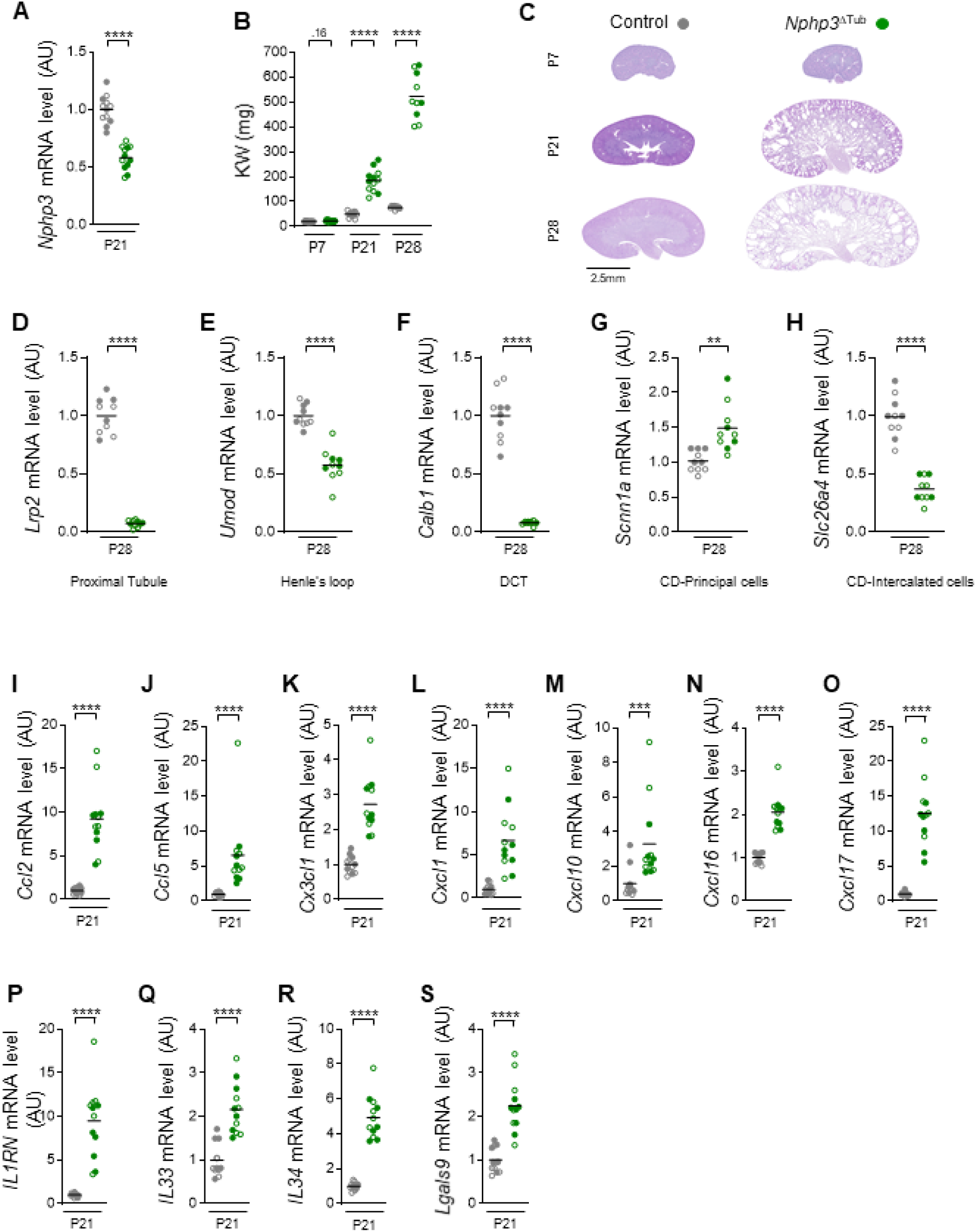
**(A)** *Nphp3* mRNA expression in kidneys from the 21 days-old control and *Nphp3*^ΔTub^ mice. **(B)** Body weight (BW) from 7, 21 and 28 days-old control and *Nphp3*^ΔTub^ mice. **(C)** Representative full kidney pictures of Periodic Acid Schiff (PAS) staining from the same 6 groups. Scale bar: 2.5mm. **(D-H)** *Lrp2* (I)*, Umod* (J)*, Calb1* (K)*, Scnn1a* (L)*, Slc26a4* (M) mRNA expression in kidneys from the 28 days-old control and *Nphp3*^ΔTub^ mice. **(I-S)** *Ccl2* (N), *Ccl5* (O), *Cx3cl1* (P), *Cxcl1* (Q), *Cxcl10* (R), *Cxcl16* (S), *Cxcl17* (T), *IL1RN* (U), *IL33* (V), *IL34* (W) and *Lgals9* (X) mRNA expression in kidneys from 21 days-old control and *Nphp3*^ΔTub^ mice. **(A-B, D-S)** Each dot represents one individual mouse. Empty circle for males, filled circles for females. Bars indicates mean. Mann-Whitney test: ** *P* < 0.01, \*\*\**P* < 0.001, \*\*\*\**P* < 0.0001. AU: arbitrary unit.

**Supplementary Figure 2.**
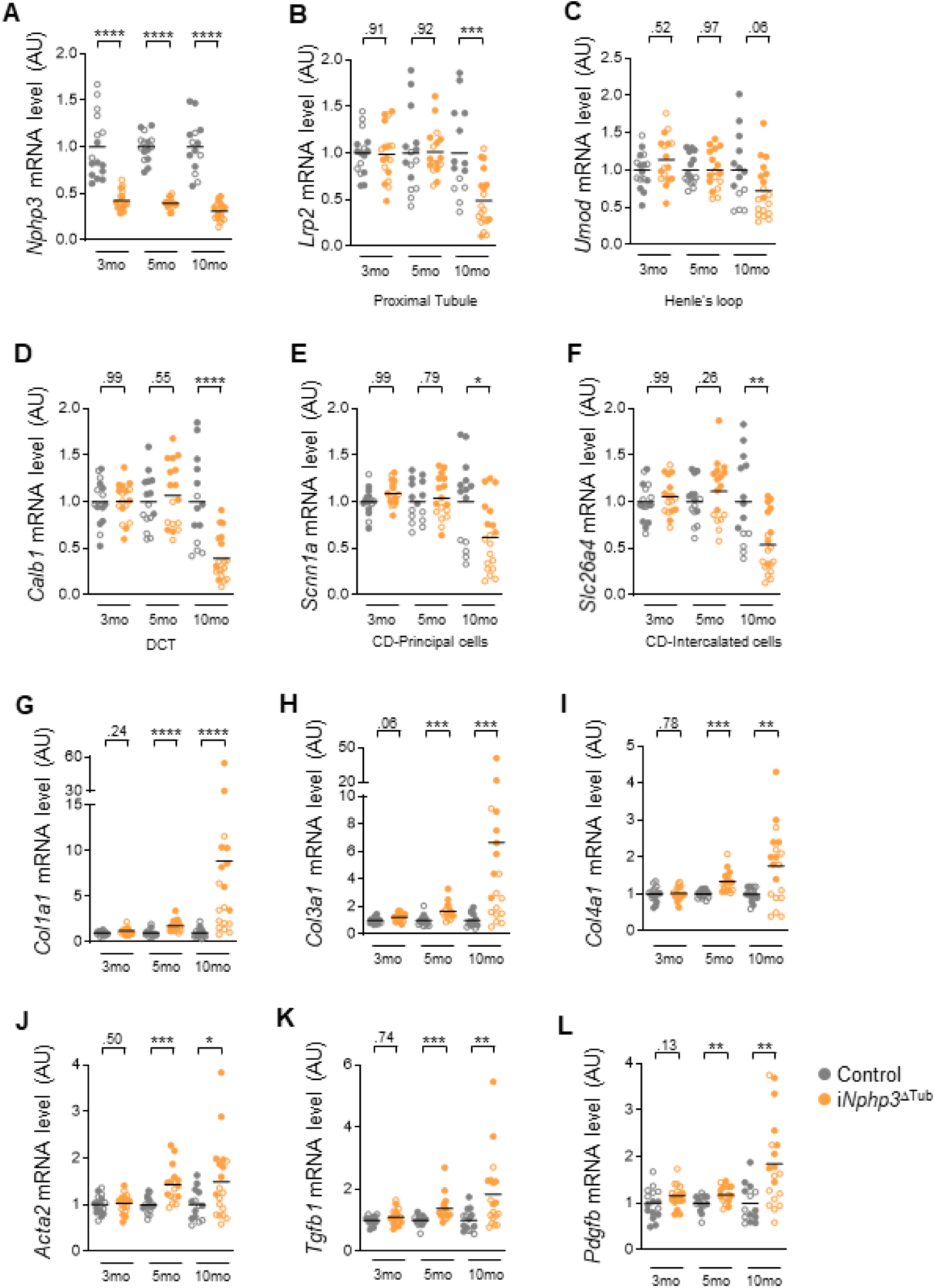
**(A-L)** *Nphp3* (A)*, Lrp2* (B)*, Umod* (C)*, Calb1* (D)*, Scnn1a* (E)*, Slc26a4* (F)*, Col1a1* (G)*, Col3a1* (H)*, Col4a1* (I)*, Acta2* (J)*, Tgfβ1* (K) *and Pdgfb* (L) mRNA expression in kidneys from 3, 5 and 10 months-old control and i*Nphp3*^ΔTub^ mice. Each dot represents one individual mouse. Empty circle for males, filled circles for females. Bars indicates mean. Mann-Whitney test: \**P* < 0.05, ** *P* < 0.01, \*\*\**P* < 0.001, \*\*\*\**P* < 0.0001. AU: arbitrary unit.

**Supplementary Figure 3.**
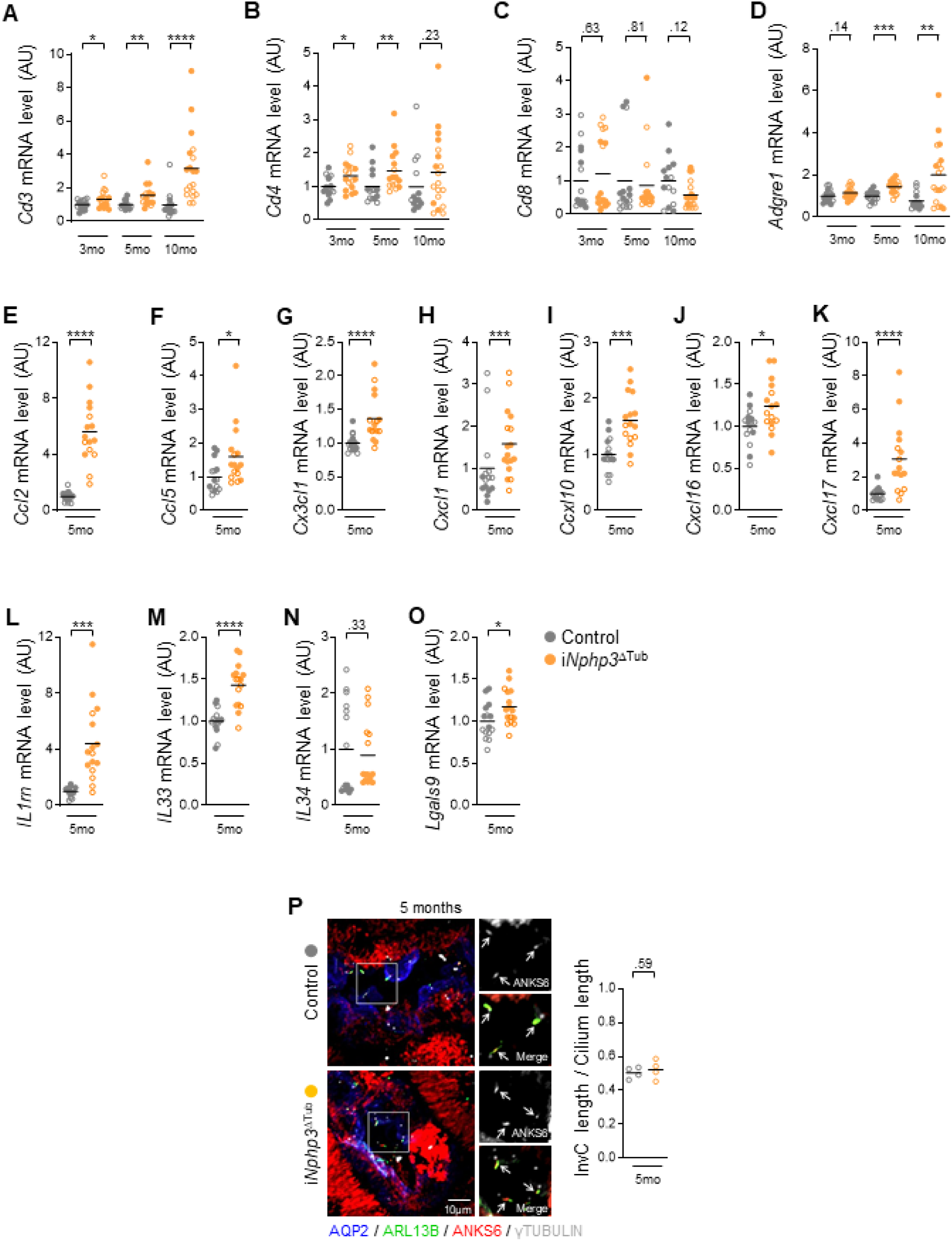
**(A-D)** *Cd3* (A)*, Cd4* (B)*, Cd8* (C) and *Adgre1* (DmRNA expression in kidneys from 3, 5 and 10 months-old control and i*Nphp3*^ΔTub^ mice. **(E-O)** *Ccl2* (E), *Ccl5* (F), *Cx3cl1* (G), *Cxcl1* (H), *Cxcl10* (I), *Cxcl16* (J), *Cxcl17* (K), *IL1RN* (L), *IL33* (M), *IL34* (N) and *Lgals9* (O) mRNA expression in kidneys from 5 months-old control and i*Nphp3*^ΔTub^ mice. **(P)** Representative images and quantification of inversin compartment length (AKSB6+, Red) normalized on primary cilia length (ARL13b+, green) in collecting ducts (CD, AQP2+, blue) of kidneys from 5 months-old control and i*Nphp3*^ΔTub^ mice. Scale bar: 10μm. Arrows indicate primary cilia. **(A-P)** Each dot represents one individual mouse. Empty circle for males, filled circles for females. Bars indicates mean. Mann-Whitney test: \**P* < 0.05, ** *P* < 0.01, \*\*\**P* < 0.001, \*\*\*\**P* < 0.0001. AU: arbitrary unit.

**Supplementary Figure 4.**
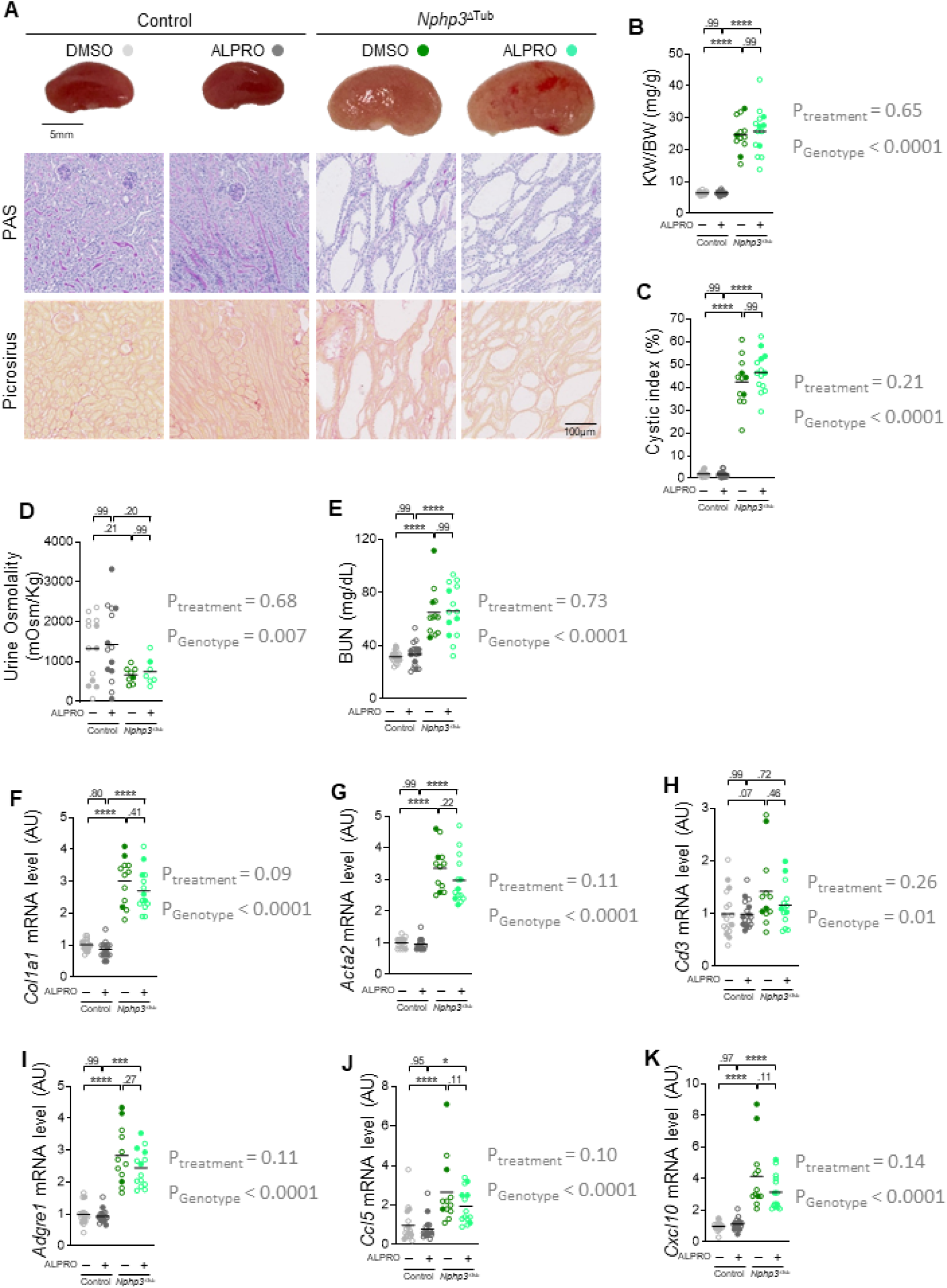
**(A)** Representative kidney pictures, Periodic Acid Schiff (PAS) and Picrosirus staining from 21 days-old (P21) control and *Nphp3*^ΔTub^ mice treated with DMSO or Alprostadil (ALPRO). Scale bar: 5mm (upper panel) and 100 µm (lower panels). **(B-E)** Kidney weight (KW) to body weight (BW) ratio, cystic index (C), Urine osmolality (D) and Blood Urea Nitrogen (BUN, E) from the same 4 mice groups. **(F-K)** *Col1a1* (F), *Acta2* (G), *Cd3* (H), *Adgre1* (I), *Ccl5* (J) and *Cxcl10* (K) mRNA expression in kidneys from the same 4 mice groups. Each dot represents one individual mouse. Empty circle for males, filled circles for females. Bars indicates mean. Two-ways anova test: \**P* < 0.05, \*\*\**P* < 0.001, \*\*\*\**P* < 0.0001. *P_treatment_* corresponding to P-value of the treatment affect and *P_Genotype_* corresponding to the P-value of the genotype affect. AU: arbitrary unit.

